# Hybrid diffuse optical monitoring of hemodynamic changes following acute cerebral ischemia in a porcine model

**DOI:** 10.1101/2020.08.07.241182

**Authors:** Detian Wang, Long Wang, Jinyu Wang, Peng Gao, Liguo Zhu, Zeren Li, Tunan Chen, Fei Li, Feng Hua

**Author notes:** Detian Wang and Long Wang contribute equally to the article.

## Abstract

Rapid screening for stroke in pre-hospital settings may improve patient outcomes by allowing early deployment of thrombolytic therapies. Near-infrared hybrid diffuse optical screening devices may fill this need. This study seeks to determine whether hybrid diffuse optical measurements can measure hemodynamic changes associated with cerebral ischemia within the first few hours of the onset of acute ischemia in a large animal model. A hybrid diffuse optical device combining of near-infrared spectroscopy (NIRS) and diffuse correlation spectroscopy (DCS) was fabricated to measure total hemoglobin concentration (HbT), tissue oxygen saturation (StO_2_) and blood flow index (BFI). Cerebral ischemia was induced by ligation of the bilateral common carotid arteries (CCA) in five miniature pigs. Additionally, a fatal stroke was induced in two pigs by injecting 5ml air emboli into both CCA. Cerebral hemodynamic parameters were monitored continuously throughout the study with the hybrid optical device. Relative changes BFI showed the good repeatability both of the ligation and fatal stroke experiments. During bilateral CCA ligation, the BFI decreased by up to about 66% of baseline values; during the fatal stroke experiment, the BFI decreased by up to about 95%, with a temporal resolution of 20 seconds. To the best of our knowledge, there are not existing methods which can measure the cerebral ischemia within the first few hours of the onset noninvasively. The MRI scanning was conducted at 24 h post injury. However, the images showed no abnormality. The results show the hybrid diffuse optical method can immediately measure the hemodynamic changes of miniature pigs in the first few hours of each single cerebral ischemia onset, and the BFI may be the promising biomarker to distinguish the cerebral ischemia and cerebral death.

## 1. Introduction

Stroke is one of the leading causes of death and disability worldwide. The only approved treatment for ischemic stroke is recanalisation of occluded arteries by thrombolysis within the very first 6 hours post infarct [1–3]. However, in clinical practice implementation of recanalising therapy within this narrow therapeutic window is difficult because neurological examination and imaging are needed. The establishment of mobile stroke units with x-ray computed tomography has the potential to substantially improve outcomes [4, 6]. However, these units require enormous financial and personnel resources to operate [7]. Therefore, less resource intensity and compact devices are disired. Near-infrared light (NIR) based devices may overcome this challenge, as these devices are sensitive to hemodynamic parameters, safe, noninvasive, cheap and portable. [8] Previous animal studies utilizing NIR techniques have focused on rat [9–16] or newborn piglet [17–20]. However, the cranial geometry of these models is significantly different from the adult human head in absolute size. Human studies have also been performed with a variety of NIRS techniques [21–33] but these studies are ethically and practically limited. In this report, we focus on monitoring the evolving injury in the first hours following an ischemic insult in a large animal porcine model.

Specifically, we utilize a hybrid diffuse optical device, which combines near-infrared spectroscopy (NIRS) and diffuse correlation spectroscopy (DCS), to measure total hemoglobin concentration (HbT), tissue oxygen saturation (StO_2_) and blood flow index (BFI). We utilized two models, both in Guangxi Bama miniature pigs: ligation of the bilateral common carotid arteries (CCA, 5 pigs) and massive air embolism (2 pigs). This result demonstrates the potential utility of DCS in monitoring ischemic events.

## 2. Method

### 2.1 Hybrid diffuse optical device

The schematic and the picture of hybrid diffuse optical device are as shown in Fig. 1. The device combines a diffuse correlation spectroscopy (DCS) and diffuse optical spectroscopy (NIRS) system [34]. The hybrid laser source consisted of one long coherent laser diode (LD, 785 nm, 90 mW, DL785-100-3O, CrystaLaser Inc., USA) and three common LD with little coherence (690 nm, 808 nm, and 830 nm with optical power levels of 50 mW, 100 mW, and 100 mW respectively, WavespectrumLaser, China). The sources were time-multiplexed to illuminate a single multi-mode fiber, integrated into an optical probe. The custom built optical probe consisted of a multi-mode source fiber (200μm/0.22NA, Nufern, USA), two single-mode (4.4μm/0.13NA, 780-HP, Nufern, USA) detection fibers with the source-detector separations of 1 cm and 2cm respectively. The light collected by the probe was detected by two single-photon avalanche photodiode detectors (SPCM-AQRH-W4, Excelitas, Canada).

**Fig.1.**
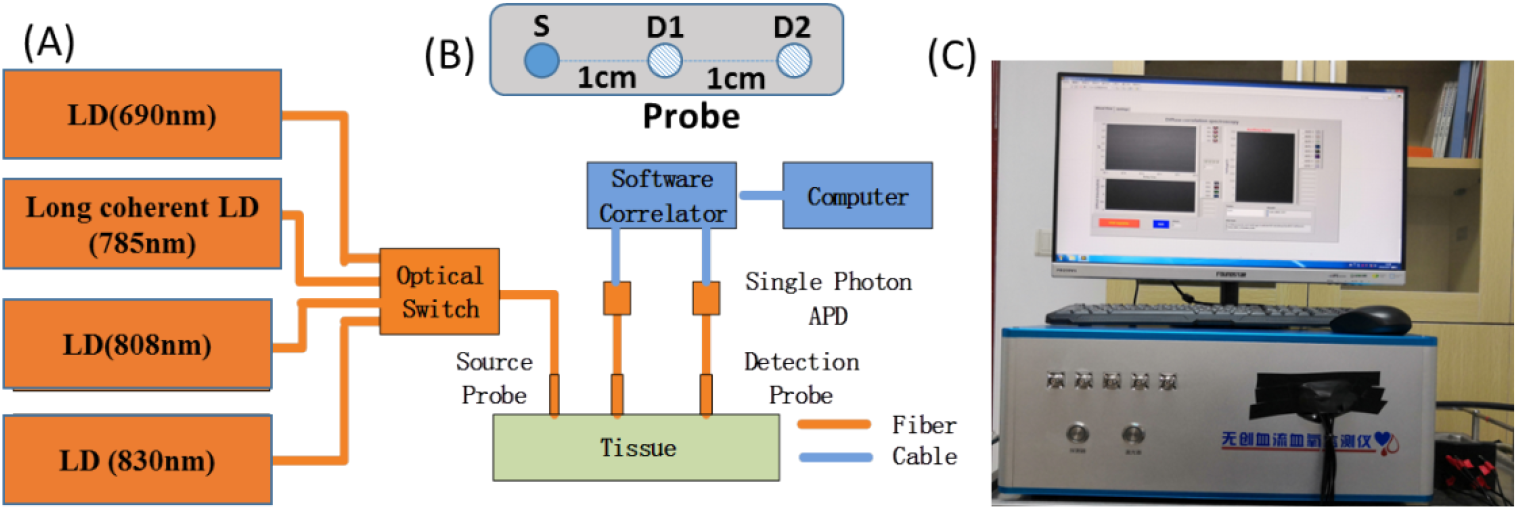
Schematic and photo of the hybrid optical device. LD: laser diode; APD: avalanche photodiode.

A custom software correlator sampled the photon arrival times for each detector and calculated both the optical intensity and the temporal autocorrelation function of the detected photons [34–36]. The BFI was derived from the decay constant of the intensity autocorrelation function (*g*_*2*_) at 785nm at 2cm, utilizing a semi-infinite solution to the photon correlation diffusion equation [37]. The HbT and StO_2_ were derived from effective attenuation coefficient of optical intensities of all four wavelengths at separations of 1 cm, and 2cm. Data were acquired at 20 Hz; each wavelength was illuminated for 5 seconds. Each multi-wavelength measurement took 20 seconds.

The absorption coefficient and the reduced scattering coefficient were assumed to be 0.1cm^−1^ and 8cm^−1^ to derive the BFI for all the pigs. As in our previous study [34], StO_2_ and HbT were derived by spatially resolved spectroscopy [38, 39]. From these, the derived variables rBFI (BFI/BFI_baseline_), dStO_2_=StO_2_-StO_2baseline_, and dHbT=HbT-Hbt_baseline_ were calculated.

Through comparing the derived BFI, StO_2_ and HbT, the changes of the derived BFI showed the better repeatability for both of the bilateral CCA ligation experiment and fatal stroke experiment than that of StO_2_ and HbT. The StO_2_ showed a decrease when the bilateral CCA were ligated or the pigs were dead, although there was a large range of values. Derived HbT changed over a broad range.

### 2.2 Experimental protocol

All animal procedures were approved by the Army Medical University. Cerebral ischemia was induced in five miniature pigs (Guangxi Bama, 25-30 kg, ages 25-34 weeks) by the ligation of bilateral CCA. A massive stroke was induced in 2 further pigs by injecting 5mL air emboli bilaterally into the CCA. Each pig was anesthetized by inhaling isoflurane (3 mL/min). After adequate anesthesia and analgesia were achieved, the pig was fixed on the operating table and the left scalp was retracted to minimize the impact of superficial tissues on the measurement. The probe was secured on the top of left skull with a bandage (position indicated by the dashed white line in Fig. 2). The bilateral CCA were exposed and a suture placed around each artery. At this moment, the pig was kept side-lying position and the optical measurement was performed for about five to ten minutes for baseline period. Then the bilateral CCA were ligated by tightening the sutures or 5mL air emboli were injected without moving the head. The hemodynamic changes were monitored for thirty to fifty minutes following induction of ischemia. A black cloth was used to cover the pigs to minimize the influence of external light. After the optical measurement, the wound was sterilized and sutured. Following recovery from anesthesia, the behavior of each pig was observed. At 24 h post injury, the extent of brain ischemia was assessed with MRI (3.0T, MAGNETOM Spectra, Siemens). Note that the bilateral CCA of five miniature pigs were always ligated through the whole experiments. After the experiments, all pigs were sacrificed by inhaling isoflurane (6 mL/min). The hemodynamic parameters between normal status and ischemic status were compared with each other. The normal status was from baseline measurement. And the ischemic status was from the last ten minutes of ischemic period measurement.

**Fig.2.**
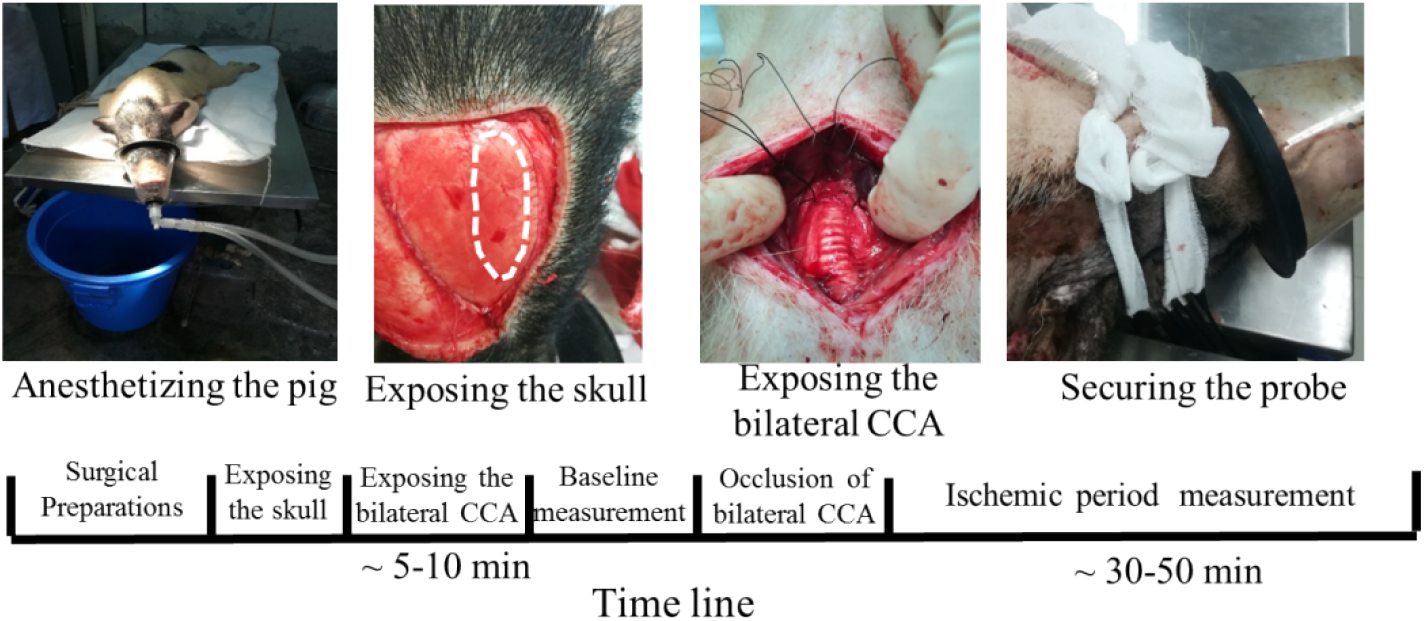
The illustrated experimental protocol.

## 3. Results

### 3.1 The bilateral CCA ligation experiment

Figure 3 shows the changes in the optical intensities and the calculated effective attenuation coefficient (*μ*_*eff*_) for all four wavelengths(690nm, 785nm, 808nm, 830nm) of an exemplar miniature pig when the bilateral CCA are occluded. The mean of optical intensities at 1cm is about 279kcps(kilo counts per second), 893kcps, 1273kcps, 1378kcps respectively, and the mean of optical intensities at 2cm are 4.2kcps, 23kcps, 33kcps, 34kcps respectively. The mean *μ*_*eff*_ are calculated as 2.85cm^−1^, 2.28 cm^−1^, 2.27 cm^−1^, 2.32 cm^−1^ respectively.

**Fig. 3.**
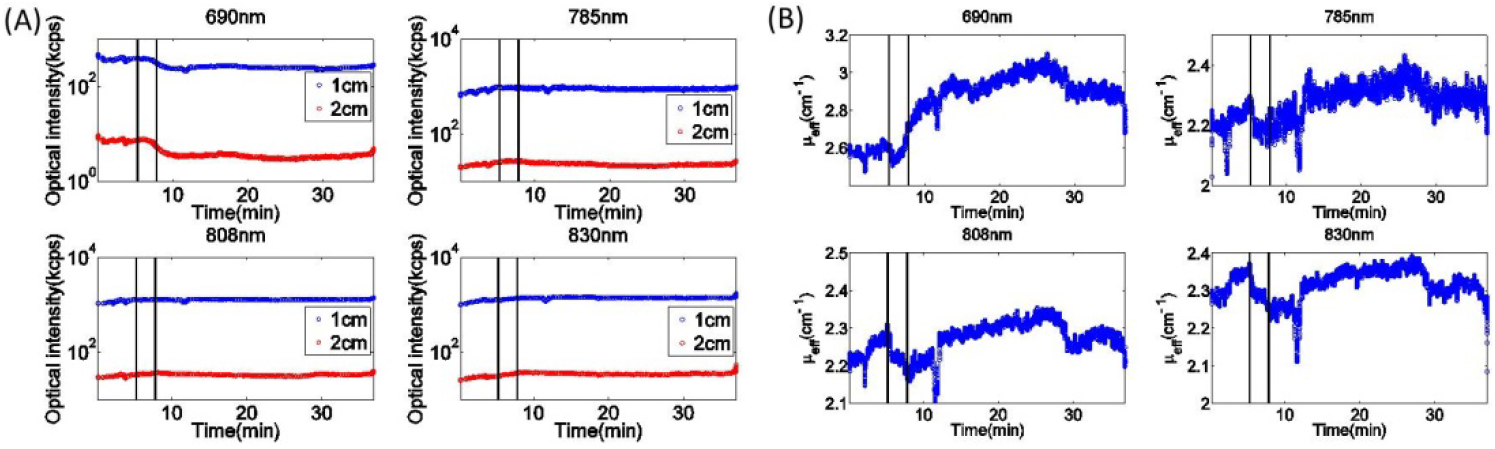
The changes in optical intensities and calculated *μ*_*eff*_ of an exemplar occlusion miniature pig

Through the effective attenuation for all four wavelengths, the StO_2_ and HbT were derived as shown in Fig. 4. The StO_2_ decreases from 60% to 46%. However, the value of HbT is quite noisy.

**Fig. 4.**
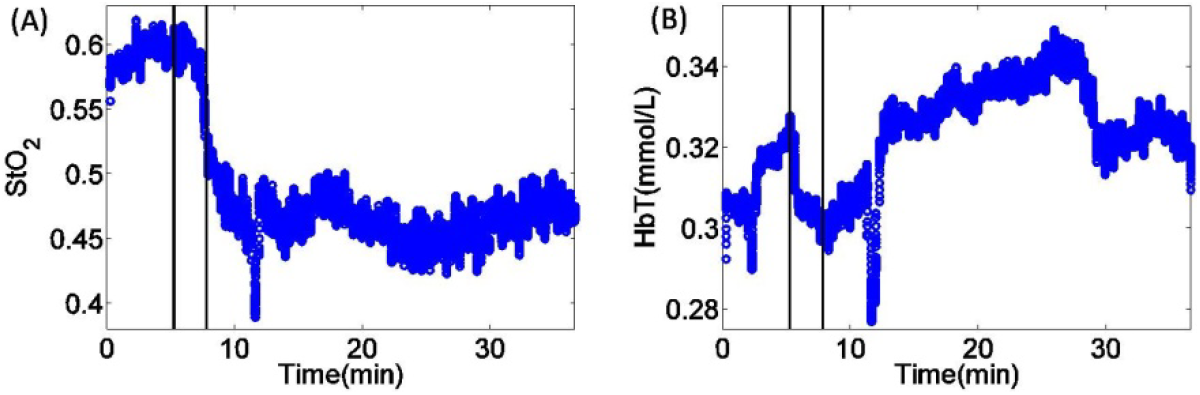
The changes in StO_2_ and HbT of an exemplar occlusion miniature pig

We also shows a representative comparison of g2 between normal and ischemic (bilateral ligated CCA) states, as shown in Fig. 5(A). As expected, the decay constant of g_2_ (BFI) becomes smaller (slower) after ligation. The BFI shows an immediate decrease up to 67% when bilateral CCA ligation is conducted, then keeps constant in thirty minutes. Fig. 3(B) shows a time course of the fitted BFI throughout the experiment. Note the rapid decrease in the BFI during the ligation (vertical black lines).

**Fig.5.**
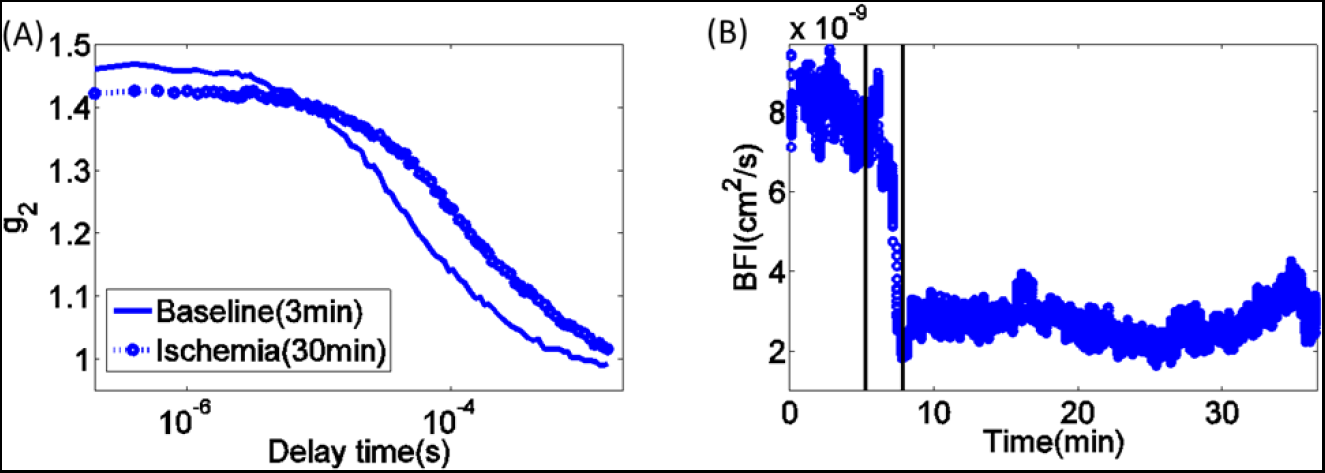
(A) The comparison of g_2_ between baseline and ischemia. (B) The BFI versus time through the bilateral CCA ligation experiment. Vertical black lines denote the period of ligation.

The BFI, StO_2_, and HbT for the five miniature pigs are as shown in Fig. 6. The results clearly shows the derived BFI and StO_2_ of all pigs decreased when the bilateral CCA were ligated. However, the decreases of StO_2_ was a large range of values. Derived HbT changed over a broad range.

**Fig. 6.**
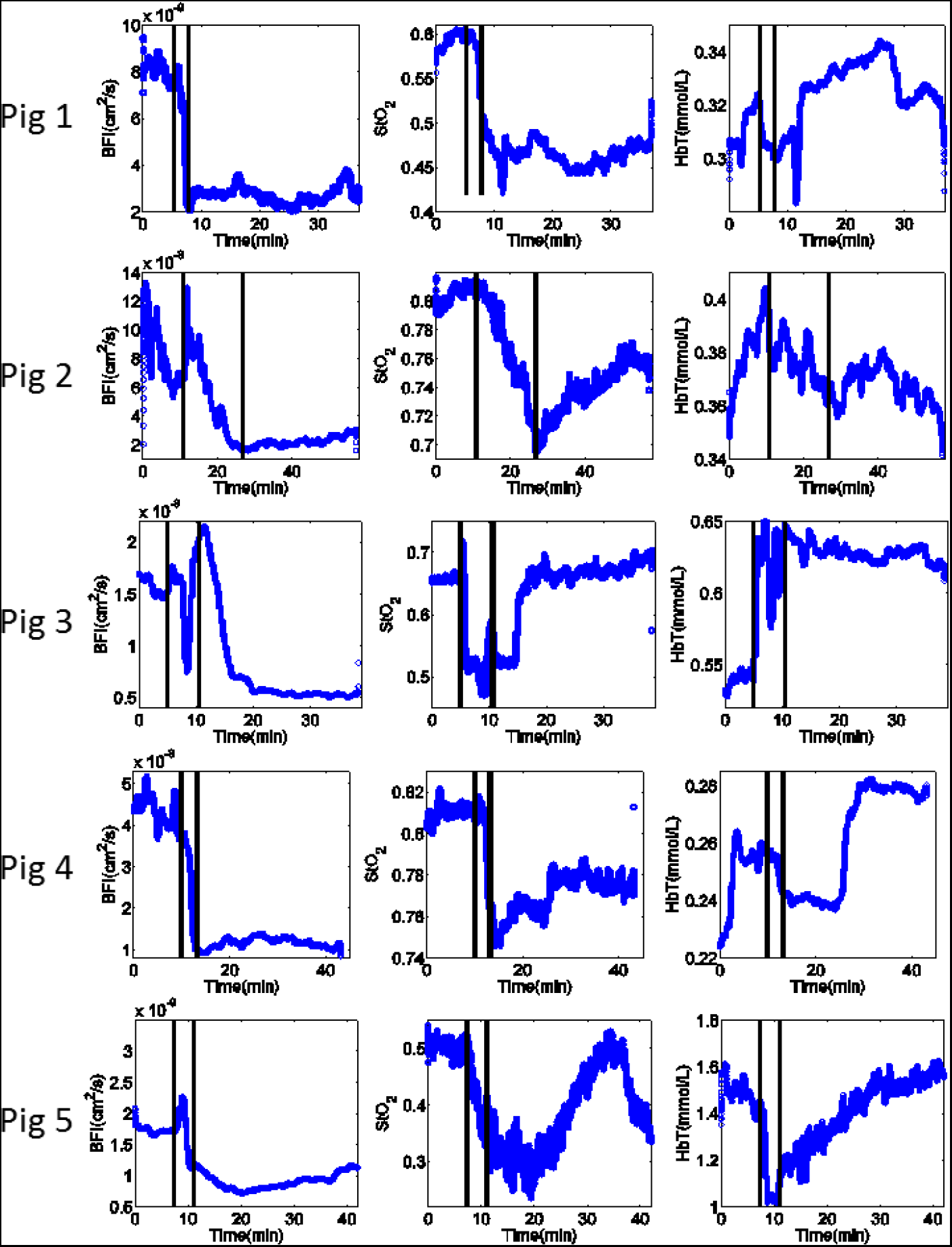
The hemodynamic changes of BFI, StO_2_ and HbT versus time for the five miniature pigs

### 3.2 The air embolism experiment

In figure 7, we show a representative case example of the air emboli experiment. Fig. 7(A) shows the comparison of g_2_ between baseline and 45 min post embolis (vertical lines, Figure 5(B)), after cardiac death. As expected, decay of g_2_ is at a much longer time scale after cardiac death. A cardiac massage was performed in this animal following cardiac death indicated by shaded area. Both pigs expired after the emboli experiment. The remnant post-mortem BFI (~5%) is the so-called biozero, due to the Brownian motion of tissue [20]. The variation in baseline BFI as shown in Fig. 7(B) was due to breathing motion artifact due to imperfect anesthesia.

**Fig.7.**
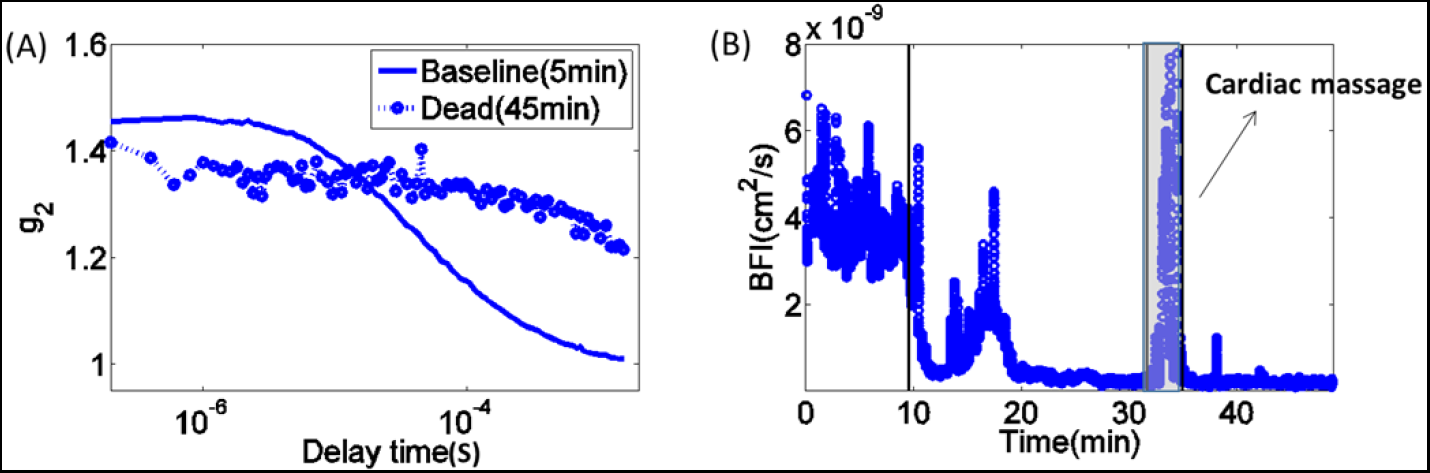
(A) The comparison of g_2_ between normal status and dead status. (B) The BFI versus time through the fatal stroke experiment.

The changes in BFI, StO_2_, and HbT versus time for the two dead pigs are as shown in Fig. 8. The results clearly show the BFI and StO_2_ had decreases after the operation.

**Fig. 8.**
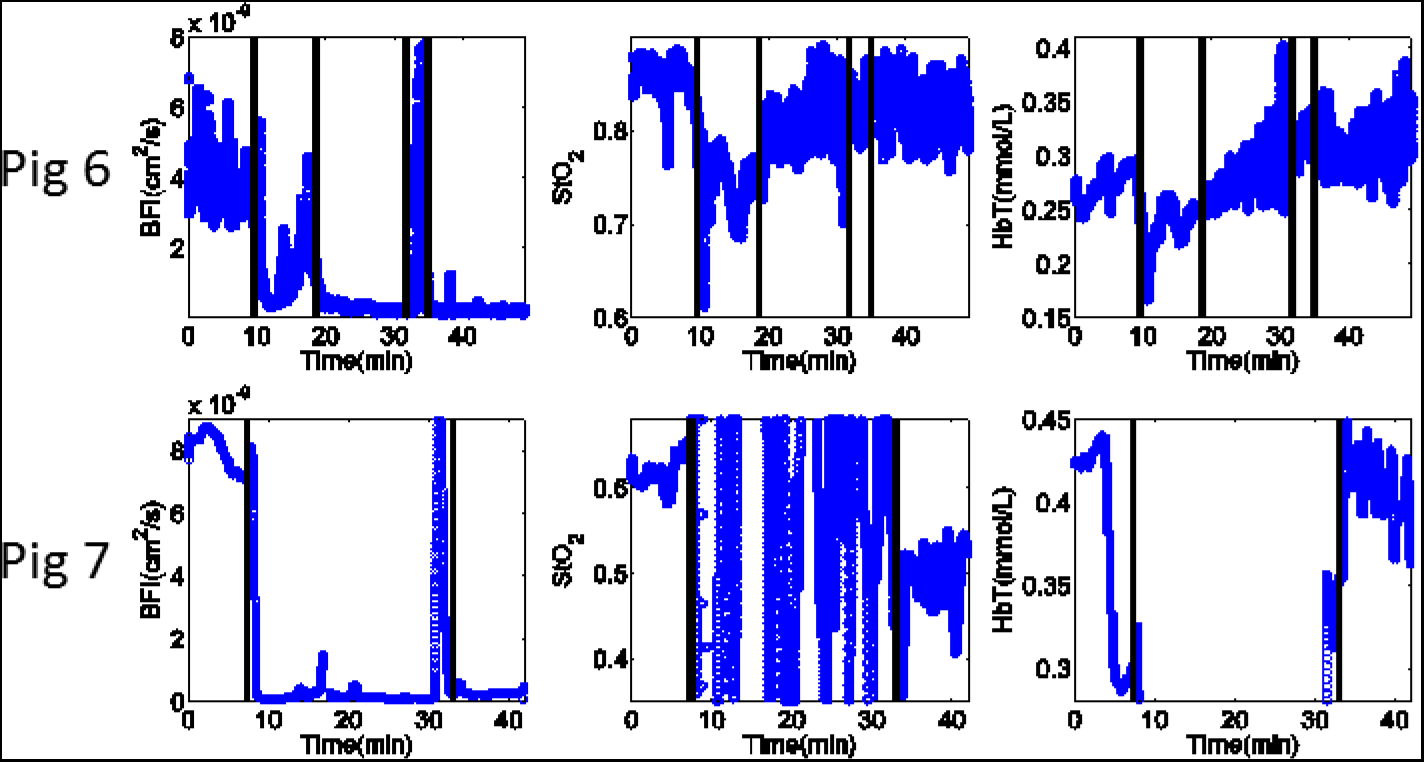
The changes in BFI, StO_2_, and HbT versus time for the dead pigs

The relative blood flow index (rBFI=BFI/BFI_baseline_) during ischemia and cardiac cessation is summarized Table 1. These data were derived from the last ten minutes of the experiment. The mean rBFI during bilateral CCA ligation is 34% and the mean rBFI following air emboli and cardiac arrest 5%.

**Table 1.** The relative changes of BFI in ischemic status and dead status.

| Operational type | Ligation of bilateral CCA |  |  |  |  | Air Embolis |  |
| --- | --- | --- | --- | --- | --- | --- | --- |
| Pig number | 1 | 2 | 3 | 4 | 5 | 6 | 7 |
| rBFI | 0.33 | 0.34 | 0.46 | 0.25 | 0.34 | 0.03 | 0.06 |

The statistic comparison of absolute BFI, StO_2_ and HbT between the normal status and the ischemic status of the ligation experiment were as shown in the Fig. 9. The baseline BFI, StO_2_ and HbT are (2.94±3.16)×10^−8^ cm^2^/s, 68%±14% and 0.59±0.51mmol/L respectively, and the BFI, StO_2_ and HbT in ischemic status are (0.93±1.06)×10^−8^cm^2^/s, 60%±20% and 0.64±0.56 mmol/L respectively. The results presented demonstrate DCS can detect the blood flow reductions due to dramatic cerebral events such as carotid ligation and the inducement of fatal strokes using air emboli. BFI The NIRS data is less conclusive though StO_2_ decreases are generally seen, while HbT changes are variable.

**Fig.9.**
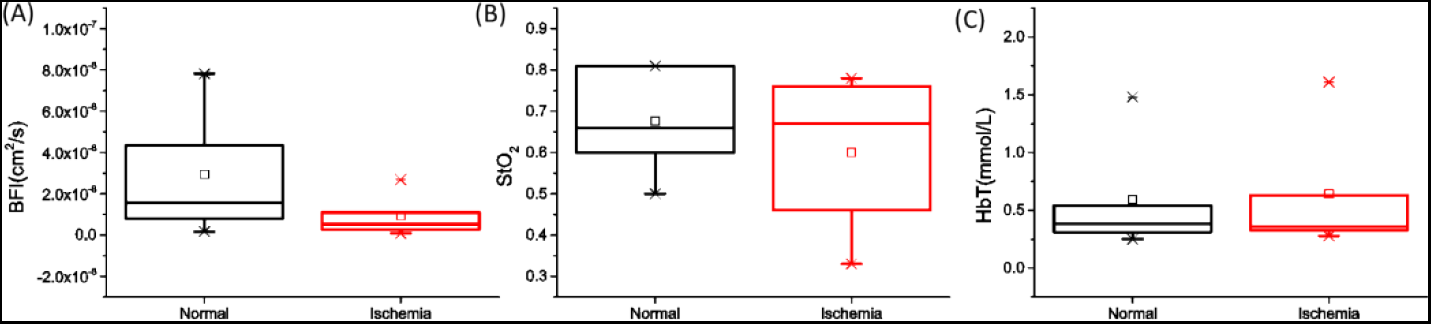
Comparison of absolute BFI, StO_2_ and HbT between the normal and the ischemic status of the ligation experiment.

The brain ischemia was assessed after 24 h with MRI as shown in Fig. 10, however no visible ischemia occurred when the when bilateral CCA ligation was conducted.

**Fig.9.**
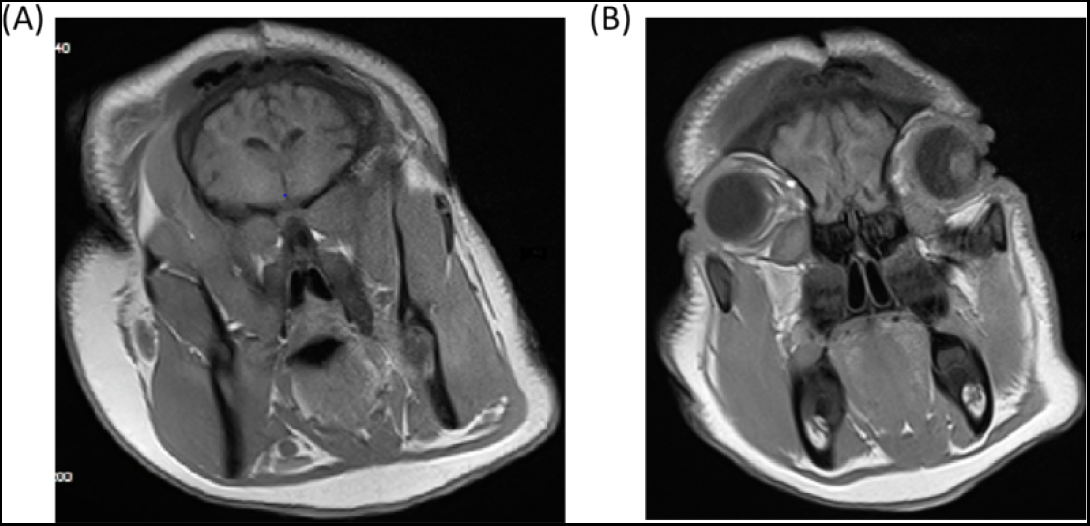
The exemplar MRI results of the miniature pig’s brain when the bilateral CCA ligation had been conducted for 24 hours

## 4. Discussion

The hybrid device has alreadly been validated to measure BFI and HbT through serveral experiments [40–49]. In this study, we investigated that whether hybrid diffuse optical measurements can measure hemodynamic changes associated with cerebral ischemia within the first few hours of the onset of acute ischemia in a large animal model. The results presented demonstrate DCS BFI can detect the blood flow reductions due to dramatic cerebral events such as carotid ligation and the inducement of fatal strokes using air emboli. The NIRS data is less conclusive though StO_2_ decreases are generally seen, while HbT changes are variable. The reason is that the NIRS data is more suffer from extraneous light and motion artifact. The DCS BFI is derived from the normalized correlation value of optical intensity, while the NIRS data is derived from the optical intensity. In the study, it’s difficult to prevent from motion artfact because miniature pig’s breath movement. Moreover, the head posture of miniature pig would be moved slightly through operation or examination. The StO_2_ is more robust that HbT, since the StO_2_ is derived from the proportion changes of spectral absorption. The 0.64 mmol/L HbT implies about 25% of tissue volume is occupied by blood since whole blood hemoglobin is around 2.3 mmol/L that seems unrealistically high. This is related to blood infiltration under the probe. Therefore, although the combine of spatially resolved spectroscopy and diffuse correlation spectroscopy has the ability to derive the absolute StO_2_, HbT and BFI, the absolute values derived from this study are questionable. But relative changes should have significant reference for such study. NIR light is sensitive to tissue changes approximately one third to half of the source-detector distance deep [50–52]. In our study, we were therefore sensitive to ~0.6-1cm deep in tissue from the 2cm separation. The results show the blood flow index decrease up to 66% which is reasonable since two thirds of the arteries which provide the blood for the brain are occluded. Unfortunately, the MRI of head of all five pigs after the bilateral CCA ligation experiment didn’t show any ischemia. Potential explanations include the difference between the techniques and the difference in time between the measurements. We did observe abnormal behavior (inability to stand steadily or walk in a straight line) in one pig, suggesting that our ischemic intervention had significant consequences.

Guangxi Bama miniature pigs are a useful animal model for ischemic stroke as both of their head size (~4cm×4cm) and skull thickness(~1cm) are close to the human head. In this experiment, we retracted the pig’s scalp(~8mm), as this is substantially thicker than that of a human and the skin-fold of scalp prevents well security of probe. The porcine cerebral vasculature is complex and includes a rete mirabile, precluding easy development of a middle cerebral artery occlusion model (MCAO) [53, 54]. As an alternative, we bilaterally ligated the CCA to produce general cerebral ischemia. These are two of the major arteries feeding the brain in the pig. To the best of our knowledge, it is the first time to investigate the feasibility of monitoring the hemodynamic changes in such acute phase of cerebral ischemia by NIR techniques on such big animal. Unfortunately the removal of the skin prior to attaching the probe breaks the connection with human studies and thus the removal of systemic physiology contamination is unrealistic and does not permit proving the utility of the technology in human studies. The most common ischemic stroke is the middle cerebral artery occlusion (MCAO). The middle cerebral artery (MCA) is the largest cerebral artery and supplies most of the outer convex brain surface, nearly all the basal ganglia, and the posterior and anterior internal capsules. Therefore the occlusion of MCA will cause the great hemodynamic changes of blood in the cortex. Since the miniature pig has comparative size of head (~4cm×4cm) and thickness of skull(~1cm) to the adult human, our results suggest that it is possible to measure hemodynamic changes of stroke in the first few hours after the onset in adult human, especially the massive MCAO stroke. This has recently been demonstrated in the therapeutic phase by Delgado-Mederos et al. during rtPA infusion [27, 30] and by Forti et al. during mechanical thrombectomy. [33]

Translation of our findings to humans requires significant additional work. Firstly, the MCAO stroke model is difficult to produce in the miniature pig; development of such a model is highly desirable. Secondly, the scalp was taken off in this study, which increases our sensitivity to cerebral tissue. Potential techniques to enhance cerebral sensitivity of these measurements include multi-parallel detection [55], few-mode fiber detection [56], multi-mode interferometric method [57], and time-domain method [58–60]. Thirdly, the deviations across the population of the absolute BFI, StO_2_ and HbT derived by the proposed hybrid diffuse optical device are quite large. This deviations may be from the inter subject variation from assumed optical properties or due to measurement accuracy. Improved measurement and analysis techniques, e.g., modelling light transport utilizing the actual head structure [61–63] will reduce these variations.

## 5. Conclusion

Through the miniature pigs experiment, we have demonstrated that the hybrid diffuse optical method can immediately measure the hemodynamic changes of miniature pigs in the first few hours following onset of cerebral ischemia. BFI may be the promising biomarker to distinguish the cerebral ischemia. Relative changes BFI showed the good repeatability for both of the ligation and fatal stroke experiments. During bilateral CCA ligation, the BFI decreased by up to about 66% of baseline values; during the fatal stroke experiment, the BFI decreased by up to about 95%, with a temporal resolution of 20 seconds.

## 7. Acknowledgments

Thanks for the discussion with David R. Busch in University of Texas Southwestern. This work is funded by the National Key Technology R&D Program of China (2017YFC1200400, 2014BA104B01, 2014BA104B05, 2015BA101B01), the Army Medical University (SWH2018QNWQ-05) and the National Natural Science Foundation of China (11604316).

